# CheckV: assessing the quality of metagenome-assembled viral genomes

**DOI:** 10.1101/2020.05.06.081778

**Authors:** Stephen Nayfach, Antonio Pedro Camargo, Emiley Eloe-Fadrosh, Simon Roux, Nikos Kyrpides

## Abstract

Over the last several years, metagenomics has enabled the assembly of millions of new viral sequences that have vastly expanded our knowledge of Earth’s viral diversity. However, these sequences range from small fragments to complete genomes and no tools currently exist for estimating their quality. To address this problem, we developed CheckV, which is an automated pipeline for estimating the completeness of viral genomes as well as the identification and removal of non-viral regions found on integrated proviruses. After validating the approach on mock datasets, CheckV was applied to large and diverse viral genome collections, including IMG/VR and the Global Ocean Virome, revealing that the majority of viral sequences were small fragments, with just 3.6% classified as high-quality (i.e. > 90% completeness) or complete genomes. Additionally, we found that removal of host contamination significantly improved identification of auxiliary metabolic genes and interpretation of viral-encoded functions. We expect CheckV will be broadly useful for all researchers studying and reporting viral genomes assembled from metagenomes. CheckV is freely available at: http://bitbucket.org/berkeleylab/CheckV.

## Introduction

Viruses are the most abundant biological entity on earth, infect every domain of life, and are broadly recognized as key regulators of microbial communities and processes [1-4]. However, it is estimated that only a limited fraction of the viral diversity on Earth can be cultivated and studied under laboratory conditions [5]. For this reason, scientists have turned to metagenomic sequencing to recover and study the genomes of uncultivated viruses [6-8]. Typically, DNA or RNA is extracted from an environmental sample, fragmented, and then sequenced, generating millions of short reads that are assembled into contigs. Metagenomic viral contigs are then identified using computational tools and algorithms [9-11] that use a variety of viral-specific sequence features and signatures. In contrast to bacteria and archaea, most viral genomes are small enough that they can be recovered by a single metagenomic contig and do not require binning (e.g. shorter than 100 kb), with notable exceptions such as the giant viruses [12] and megaphages [13] with large genomes, or segmented viruses.

Assembly of viruses from metagenomes is challenging [14] and the completeness of assembled contigs can vary widely, ranging from short fragments to complete or near-complete genomes [15]. Small genome fragments may adversely affect downstream analyses including estimation of viral diversity, host prediction, or identification of core genes across viral taxa. Viral contigs can also be derived from integrated proviruses, in which case the viral sequence may be flanked on one or both sides by regions originating from the host genome. This type of host contamination also adversely affects downstream analyses, especially the estimation of viral genome size, characterization of viral gene content, and identification of viral-encoded metabolic genes [16].

For bacteria and archaea, genome quality can now be readily determined. The most widely adopted method, CheckM, estimates genome completeness and contamination based on the presence and copy number of widely distributed, single copy marker genes [17]. Because viruses lack widely distributed marker genes, the most commonly used approach with regard to completeness is to apply a uniform length threshold (e.g. 5 or 10 kb) and analyze all viral contigs longer than this length. However, this “one-size-fits all” approach fails to account for the large variability in viral genome sizes, which range from 2 kb in *Circoviridae* [18] up to 2.5 Mbp in *Megaviridae* [12], and thus gathers sequences representing a broad range of genome completeness. Complete, circular genomes are commonly identified from the presence of direct terminal repeats [5-7], and sometimes from mapping paired-end sequencing reads [19], but tend to be rare. VIBRANT [11] is a recently published tool that categorizes sequences into high-, medium-, or low-quality tiers based on circularity and the presence of viral hallmark proteins, but does not estimate genome completeness per se.

With regard to contamination, existing approaches either remove viral contigs containing a high fraction of microbial genes [5] or predict host-virus boundaries on proviruses [10, 11, 20, 21]. The former approach allows for a small number of microbial genes while the latter approach may misidentify the true host-virus boundary. Other approaches detect viral signatures, but do not account for the presence of microbial regions whatsoever [9]. With the diversity of available viral prediction pipelines and protocols, there is a need for a standalone tool to ensure that viral contigs are free of host contamination, and to remove it when present.

Here, we present CheckV, a new tool to automatically estimate completeness and contamination for metagenome-assembled viral genomes. By collecting an extended database of complete viral genomes from both isolates and environmental samples, CheckV was able to estimate the completeness for the vast majority of contigs in the IMG/VR database, illustrating its broad applicability to newly assembled genomes across viral taxa and Earth’s biomes. In addition, CheckV uses a new approach comparing the gene content between contiguous sliding windows along each sequence to identify putative host contamination on contig’s edges stemming from the assembly of integrated proviruses. Application to the IMG/VR database revealed that this type of contamination was rare but could easily lead to wrongful interpretation of viral genome size and viral-encoded metabolic genes.

## Results

### A framework for assessing viral genome quality

CheckV is a fully automated, command-line tool for assessing the quality of metagenome-assembled viral genomes. It is organized into three modules which identify and remove host contamination on proviruses (Figure 1A), estimate completeness for genome fragments (Figure 1B), and predict closed genomes based on terminal repeats and provirus integration sites (Figure 1C). Based on these results, the program classifies each sequence into one of five quality tiers (Figure 1D) - complete, high-quality (>90% completeness), medium-quality (50-90% completeness), low-quality (0-50% completeness), or undetermined-quality (no completeness estimate available) - which are consistent with and expand upon the MIUViG standards [15]. Because host contamination is easily removed, it is not factored into these quality tiers.

**Figure 1.**
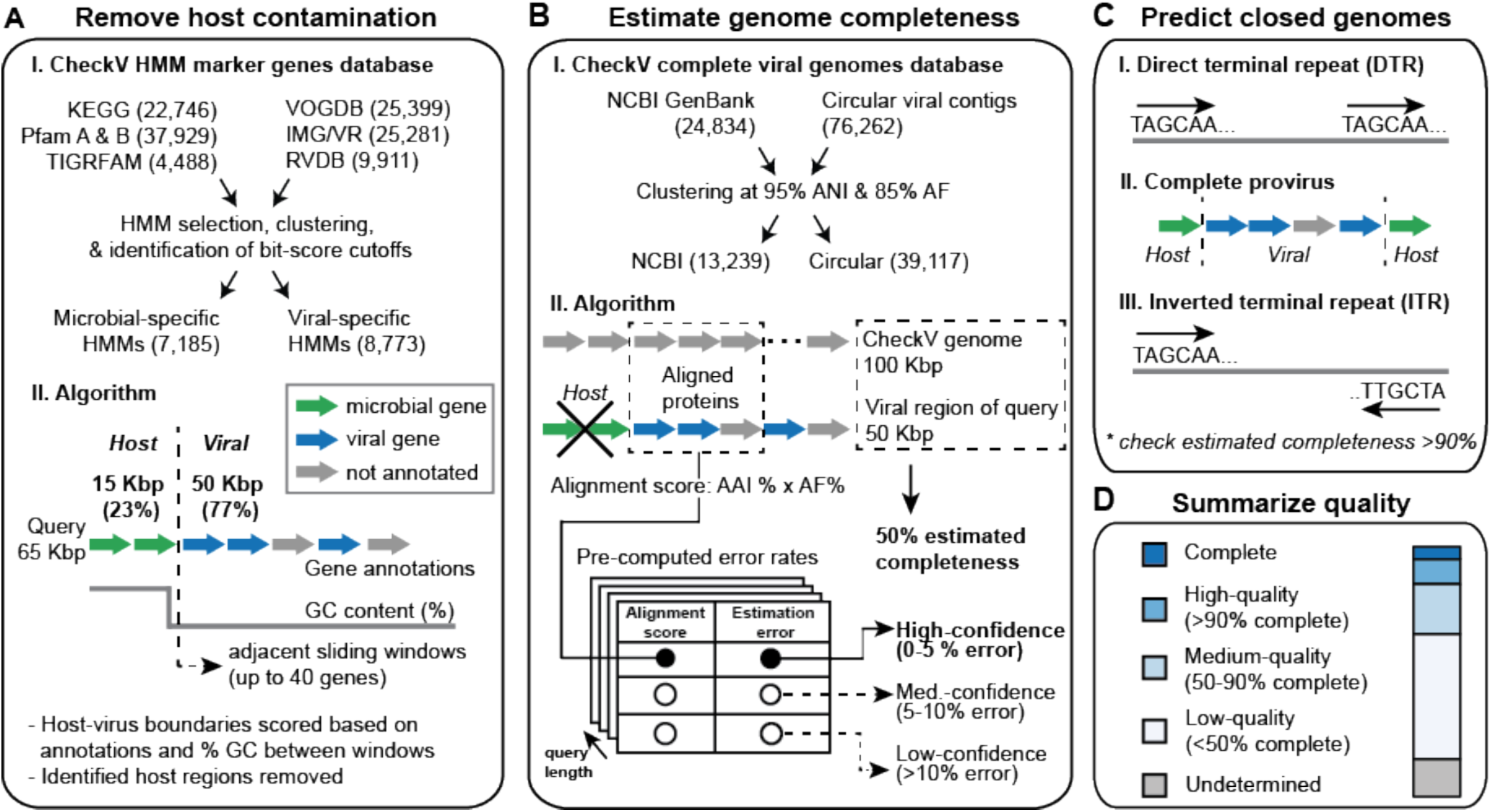
A framework for assessing the quality of metagenome-assembled viral genomes. CheckV estimates the quality of viral contigs from metagenomes in three main steps. A) First, CheckV identifies and removes non-viral regions on proviruses using an algorithm that leverages gene annotations and GC content. B) CheckV estimates the genome completeness based on AAI to a large database of complete viral genomes derived from NCBI GenBank and environmental samples and reports a confidence level for the estimate. C) Closed genomes are identified based either on direct terminal repeats, prophage integration sites, or inverted terminal repeats. When possible, these predictions are validated based on the estimated completeness obtained in B. D) Finally, sequences are assigned to one of five different quality tiers based on their estimated completeness. (ANI: average nucleotide identity; AF: alignment fraction; AAI: average amino acid identity).

In the first step, CheckV identifies and removes non-viral genes on the edges of contigs, which can occur for assembled proviruses (Figure 1A and Methods). Genes are first annotated as either viral or microbial (i.e. from bacteria or archaea) based on comparison to a large database of 15,958 profile hidden markov models (HMMs) (Figure S1 and Table S1). We selected these HMMs from seven reference databases using three main criteria: high specificity to either viral or microbial proteins, commonly occurring in either viral or microbial genomes, and non-redundant compared other HMMs. Starting at the 5’ edge of the contig, CheckV compares these gene annotations as well as GC content between a pair of sliding windows that each contain up to 40 genes. This information is then used to compute a score at each intergenic position and identify host-virus boundaries. We optimized this approach to sensitively and specifically detect flanking host regions, even those containing just a single gene.

In the second step, CheckV estimates the expected genome length of contigs based on the average amino acid identity (AAI) to a database of complete viral genomes from NCBI and environmental samples (Figure 1B and Methods). The expected genome length is then used to estimate completeness as a simple ratio of lengths. In contrast to bacteria and archaea, genome size is relatively conserved among related viruses, particularly at the family or genus ranks [15], which enables CheckV to infer the expected genome length of a new virus based on its hits to the genome database. For example, genome lengths differed by only 12.5% on average (IQR = 4.2% to 16.7%) for viruses displaying just 30-40% AAI over at least 10% of their genes (Figure S2).

Rather than set an arbitrary threshold, we empirically derived the relationship between genome similarity and genome size variation, which is stored as a lookup table in the database. Using this table, together with the observed similarity and contig length, CheckV reports the confidence of each estimate: high-confidence (0-5% error), medium-confidence (5-10% error), or low-confidence (>10% error).

Highly novel viruses can be too diverged from CheckV genomes to obtain a high- or medium-confidence completeness estimate based on AAI. In these cases, CheckV uses a more sensitive HMM-based approach. After annotation with the viral HMMs (Figure 1A), CheckV compares the length of the viral contig to the lengths of CheckV reference genomes that are annotated by the same HMMs. Using this information, CheckV is able to obtain a conservative estimate of genome completeness (i.e. maximum completeness value that can be ascertained with >95% probability).

In the last step, CheckV predicts closed genomes based on direct terminal repeats (DTRs), inverted terminal repeats (ITRs), and provirus integration sites. CheckV identifies DTRs and ITRs based on a repeated sequence of at least 20-bp at the start and end of a contig. While DTRs can play a role in genome integration [22], they largely result from assembling short reads from a circular genome [23] or a linear genome that has been circularly permuted by a replication mechanism involving a concatemer intermediary [24]. Inverted terminal repeats (ITRs) are a hallmark of transposons [25] but have also been observed in complete viral genomes [26] and phages [27]. Complete proviruses are identified by a viral region flanked by host DNA on both sides, which is a commonly used approach [10, 11, 20, 21].

### An expanded database of complete viral genomes from metagenomes

We initially formed the CheckV genome database using 24,834 viral genomes from NCBI GenBank [28] (Table S2). However, uncultivated identified viruses commonly display little to no similarity to reference databases [5]. To mitigate this issue and expand the diversity of the database, we used CheckV to perform a systematic search for metagenomic viral contigs with DTRs (DTR contigs) from over 14.4 billion contigs (9.7 Tb) derived from publicly available and environmentally diverse metagenomes, metatranscriptomes, and metaviromes downloaded from: IMG/M [29], MGnify [30], and recently published studies of the human microbiome [31-33] and ocean virome [6] (Figure 2 and Methods). We exclusively used DTRs to identify complete genomes as this is the most well-established approach [5-7].

**Figure 2.**
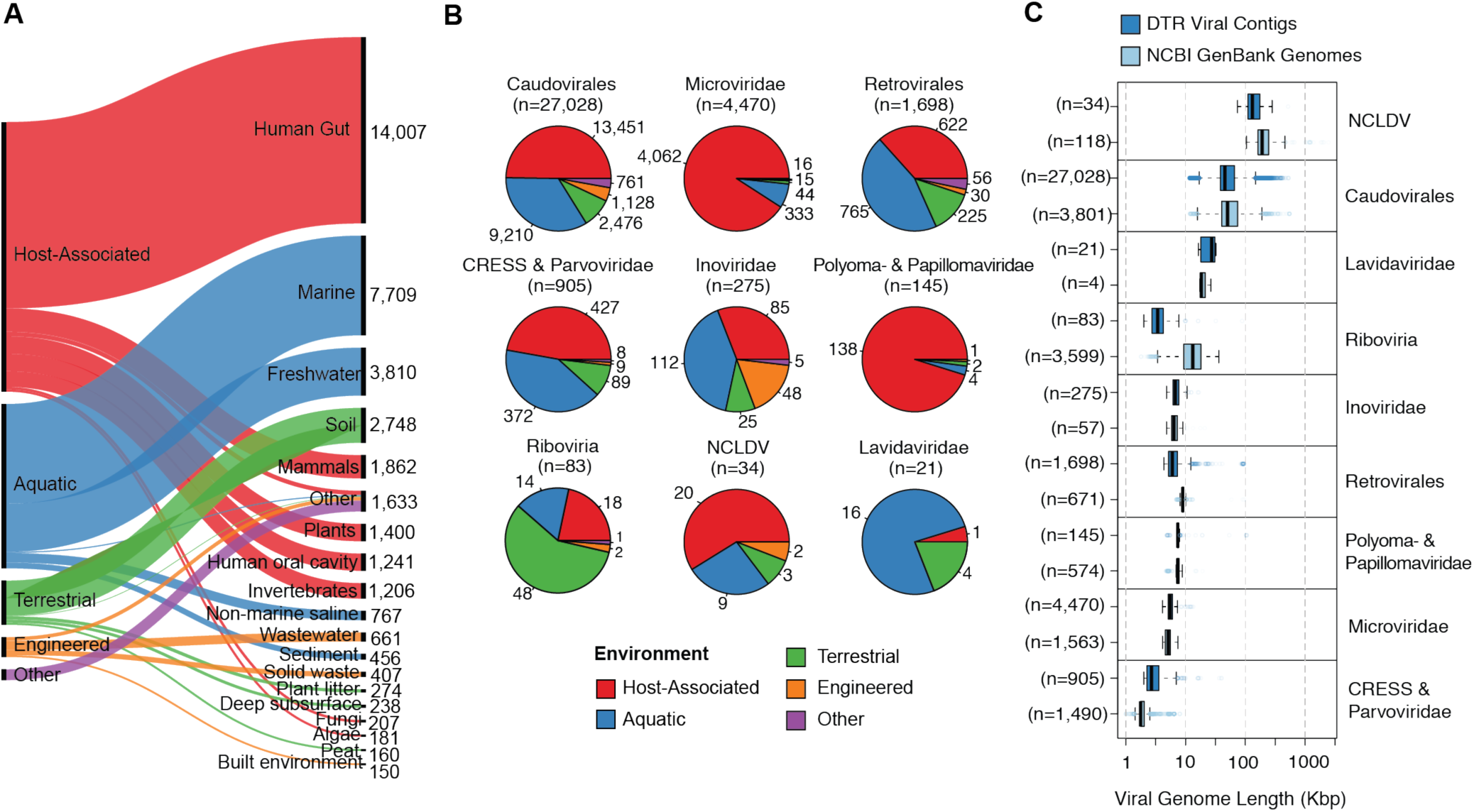
An expanded reference database of environmentally diverse genomes. 76,262 DTR contigs were identified from publicly available metagenomes, metatranscriptomes, and viromes and were clustered into 39,117 non-redundant genomes at 95% ANI. A) The non-redundant genomes are derived from diverse human-associated and environmental habitats. Habitats are based on the GOLD (Genomes OnLine Database) ontology [34] and visualization was made using RAWgraphs [35]. B) The 39,117 genomes were taxonomically annotated based on clade-specific marker genes from the VOG database. C) Comparison between GenBank genomes and DTR contigs for different viral clades.

Using this approach, we identified 76,262 DTR contigs after carefully filtering out potential false positives and verifying completeness (Figure S3 and Table S3). These were subsequently de-replicated to 39,117 sequences at 95% ANI (average nucleotide identity) over 85% of the length of both sequences (Table S4). DTR contigs were found in diverse environments including human gut (35.8%), marine (19.7%), freshwater (9.7%), and soils (7.0%) and were derived from major clades of DNA viruses, including *Caudovirales* (69.1%), *Microviridae* (11.4%), and CRESS viruses (2.3%) (Figure 2A-B). DTR contigs were also identified for RNA viruses (i.e. *Riboviria*, N=83) and retroviruses (i.e. *Retrovirales*, N=1,698), which were further confirmed through identification of marker genes (e.g. RdRp) and association to known viral families (Supplementary Information).

Next, we compared the 76,262 DTR contigs to the 24,834 GenBank references and de-replicated all sequences again at 95% ANI resulting in 52,141 clusters. Overall, the addition of DTR contigs resulted in a 3.9-fold increase in the number of clusters (Figure 1B), which was particularly pronounced for the *Caudovirales* order (7.1-fold increase) (Figure 2B). In contrast, GenBank genomes had improved representation of other viral clades, including RNA viruses from the *Riboviria* realm (Table S5). For most viral clades, the sizes of DTR contigs and GenBank genomes were consistent, indicating no systematic artifacts in our data (Figure 2B). One interesting exception was for segmented RNA viruses (*Riboviria* and *Retrovirales*), in which the DTR contigs tended to be smaller than the GenBank references, suggesting they represent a single genome segment or may only cover a subset of the diversity within these large groups.

### Accurate estimation of completeness and contamination

Having developed the CheckV pipeline and databases, we next benchmarked its accuracy. To evaluate genome completeness, we generated a mock dataset containing fragments from 382 bacteriophages not included in the CheckV database (Table S6). Using a combination of the AAI- and HMM-based approaches, CheckV estimated completeness with a median unsigned error (MUE) of only 0.91% (Figure 3A-B). Several of the genomes in the mock dataset were closely related to a CheckV reference, which may be unrealistic when analyzing real metagenomes. To simulate novel viruses, we reran the program using only reference genomes that displayed low similarity to genomes in the mock dataset (Table S7). As similarity decreased, CheckV automatically switched from using the AAI-based approach to the more sensitive, but less accurate HMM-based approach. As a result, the MUE increased from 0.91% when using the full CheckV database to 5.5% when using only highly diverged references (e.g. <30% AAI compared to the mock dataset), but enabled CheckV to estimate completeness for >96% of all fragments.

**Figure 3.**
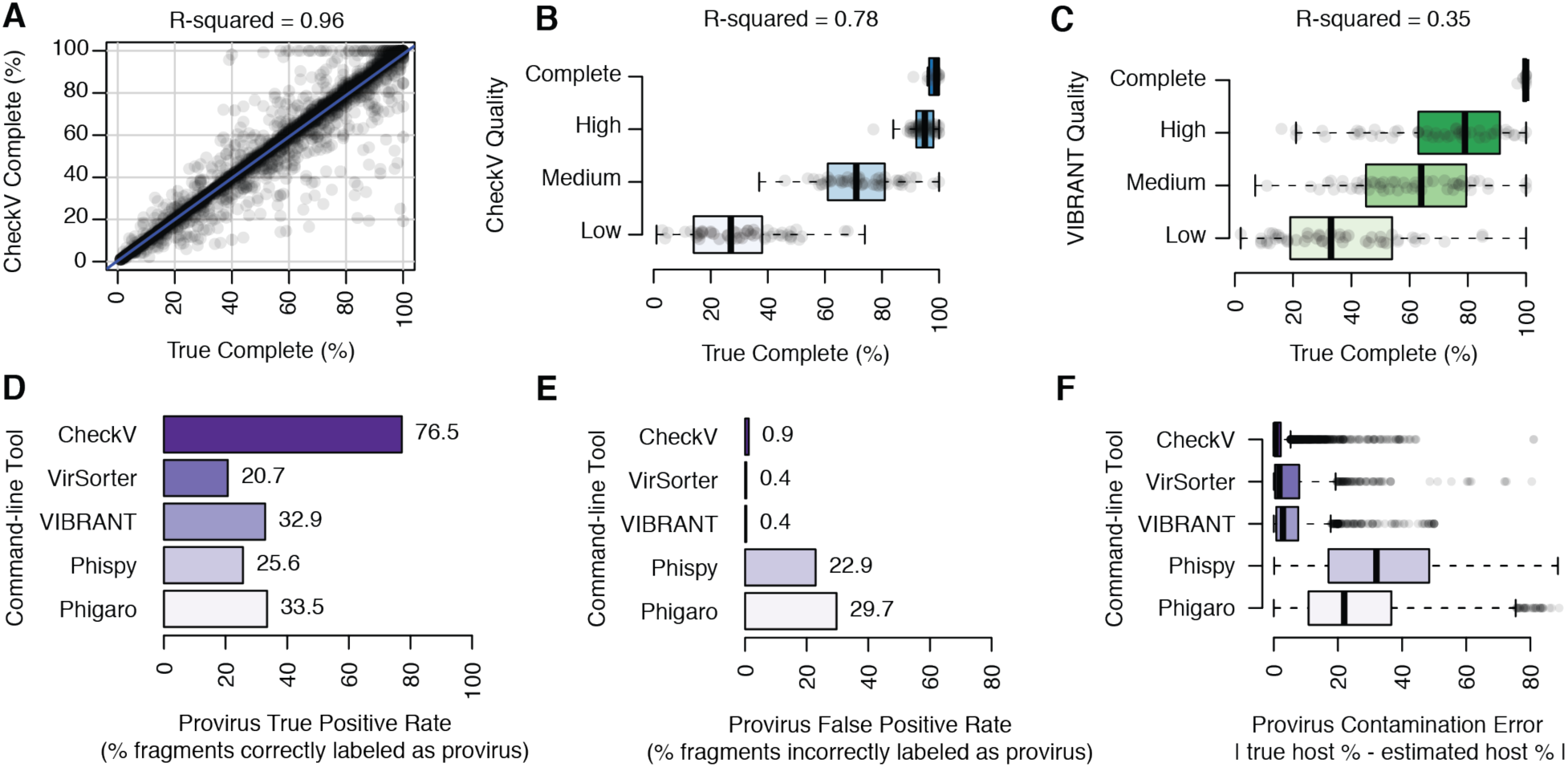
Benchmarking CheckV and comparison with existing tools. A-C) Benchmarking genome completeness on a mock dataset of bacteriophage genome fragments. A) CheckV’s estimated completeness versus true completeness. B) CheckV’s quality tiers versus true completeness. C) VIBRANT’s quality tiers versus true completeness. D-F) Benchmarking detection of host regions for CheckV and existing tools on a mock dataset of genome fragments from proviruses. Proviruses were identified as any genome fragment with a predicted viral region that covered <95% of its length. D) The percent of provirus fragments correctly identified by each tool. E) The percent of viral fragments incorrectly classified as proviruses by each tool. F) The error in estimated contamination (i.e. percent of contig length derived from host genome) for correctly identified proviruses.

For comparison, we applied VIBRANT to the mock dataset, which is currently the only available tool to assess viral genome completeness (Figure 3C). VIBRANT does not estimate completeness, per se, but it assigns fragments to four quality tiers. Compared to the true completeness values, VIBRANT’s quality tiers displayed lower correlations (R2 = 0.35) than CheckV’s estimated completeness (R2 = 0.96) or CheckV’s quality tiers (R2 = 0.78).

Next, we evaluated CheckV’s accuracy in detecting host contamination on provirus sequences (Table S8). To generate mock proviruses, we paired the 382 bacteriophages with their bacterial and archaeal hosts from the Genome Taxonomy Database [36], inserting each phage at a random location in its host genome, and extracting genome fragments of varying length (5 to 50 kb) and amount of host sequence (10 to 50%). As a negative control, we generated genome fragments that were entirely viral. Overall, CheckV correctly classified 76.5% of mock proviruses (Figure 3D) while incorrectly classifying only 0.9% of the entirely viral sequences (Figure 3E). CheckV was also able to accurately estimate the length of the host region on proviruses (Figure 3F) but was less sensitive for shorter genome fragments (Figure S4).

For comparison, we evaluated four commonly used tools for identifying host-provirus boundaries, including VIBRANT [11], VirSorter [10], PhiSpy [20], and Phigaro [21]. To enable comparability between tools, proviruses were defined as any genome fragment with a predicted viral region that covered <95% of the fragment length. Overall, none of the tools sensitively detected mock proviruses (range = 20.7 to 33.5%; Figure 3D), particularly when fragments were short, or host contamination was low (Figure S4). For example, VirSorter detected only 1.5% of proviruses with <20% contamination and only 5.1% shorter than 20 kb. This implies that microbial genes at the edges of viral contigs may be overlooked by existing tools and interpreted as viral-encoded functions. In contrast, other tools identified host-virus boundaries on entirely viral sequences (Figure 3E). For example, PhiSpy predicted non-viral regions on 22.9% of entirely viral fragments which covered 26.3% of the length of these sequences on average. This implies that truly viral regions may be discarded with existing tools and that sequences may be inadvertently classified as integrated proviruses.

Finally, we compared the computational efficiency of CheckV to existing tools. Using 16 CPUs (Intel Xeon E5-2698 v3 processors), CheckV was 1.6× to 11.6× faster than the other programs when applied to the mock dataset and required ∼2 GB of RAM. Using a single CPU CheckV was still faster than VirSorter and VIBRANT but slower compared to PhiSpy and Phigaro (Table S9).

### Using CheckV to identify high-quality genomes from metagenomes and viromes

To illustrate the type of results obtained with CheckV and its ability to scale to large datasets, we first applied it to the 735,106 contigs from the IMG/VR 2.0 database [37] (Table S10). The IMG/VR contigs were identified in assembled metagenomes using the Earth’s Virome Protocol [5] using a minimum length cutoff of 5 kb. The original samples came from many studies, the majority of which did not use size filtration to enrich for extracellular viral particles. Because of the sample characteristics and detection approach, this dataset is mostly composed of environmental dsDNA phages and contains sequences from both lysogenic and lytic viruses.

First, we used CheckV to identify three types of complete genomes from IMG/VR, including: DTR contigs (N=14,844), proviruses with 5’ and 3’ attachment sites (N=1,052), and contigs with inverted terminal repeats (ITRs; N=579). The longest DTR contig we identified was a 528,258 bp sequence from a saline lake in Antarctica estimated to be 100.0% complete and supported by paired end reads that connected contig ends. Based on gene content and phylogeny, this sequence is likely a member of one of the recently defined clades of “huge” phages [13] (Supplementary Text, Figure S5). To validate the other potentially complete genomes, we compared the contigs to CheckV’s database of complete reference genomes based on AAI, estimated completeness (medium- and high-confidence estimates only), and identified high-quality assemblies (i.e. >90% complete). We found that 90.1% of the DTR contigs with estimated completeness met the high-quality standard, compared to only 58.2% of complete proviruses and 31.8% of ITRs. In the case of proviruses, lower estimated completeness may be due to their domestication and degradation in the host genome over time [38]. These results confirm that DTRs are a good indicator of complete viral genomes most of the time [15], but suggest that greater caution is needed when interpreting other signatures.

Next, we used CheckV to estimate completeness for the entire IMG/VR dataset, including genome fragments. Using the accurate AAI-based approach, completeness could be estimated for 572,369 IMG/VR contigs (78.6%) with high- or medium-confidence, including 82.8% from host-associated, 82.0% from marine, 72.2% from freshwater, and 65.9% from soil environments. For the majority of these contigs, the best hit in the CheckV database was a DTR sequence (N=486,130, 84.9%) and was often derived from the same habitat as the IMG/VR contig (Figure S6). We next applied the HMM-based approach, which increased the percent of IMG/VR contigs with estimated completeness to 97.8%. The AAI- and HMM-based estimates of completeness were well correlated for IMG/VR contigs with both predictions (Spearman’s rho = 0.90), but the HMM-based estimates were consistently lower (94.1% of contigs), which is expected given that this approach was designed to be more conservative.

We next classified IMG/VR sequences into quality tiers according to their estimated completeness, revealing 1.9% complete, 2.5% high-, 6.5% medium-, and 86.9% low-quality sequences, with the remainder 2.2% with undetermined quality (Figure 4A). Contig sizes were strongly correlated with quality tiers, with complete genomes centered at 44 kb, which is consistent with genome sizes from the *Caudovirales* order (Figure 4B). Interestingly, aquatic samples, both marine and freshwater, seemed to contribute more partial genomes than other environments, which may reflect challenges in metagenomic assembly for these environments related to low sample biomass [39], high strain heterogeneity [40], or rare (i.e. low-coverage) viral populations. We also applied CheckV to the Global Ocean Virome (GOV) 2.0 dataset [6] (Table S11) revealing remarkably similar patterns (Figure S7). Like IMG/VR, the GOV dataset contains viral contigs that are at least 5 kb, but unlike IMG/VR, the original samples were from the open ocean and enriched for viral particles prior to sequencing.

**Figure 4:**
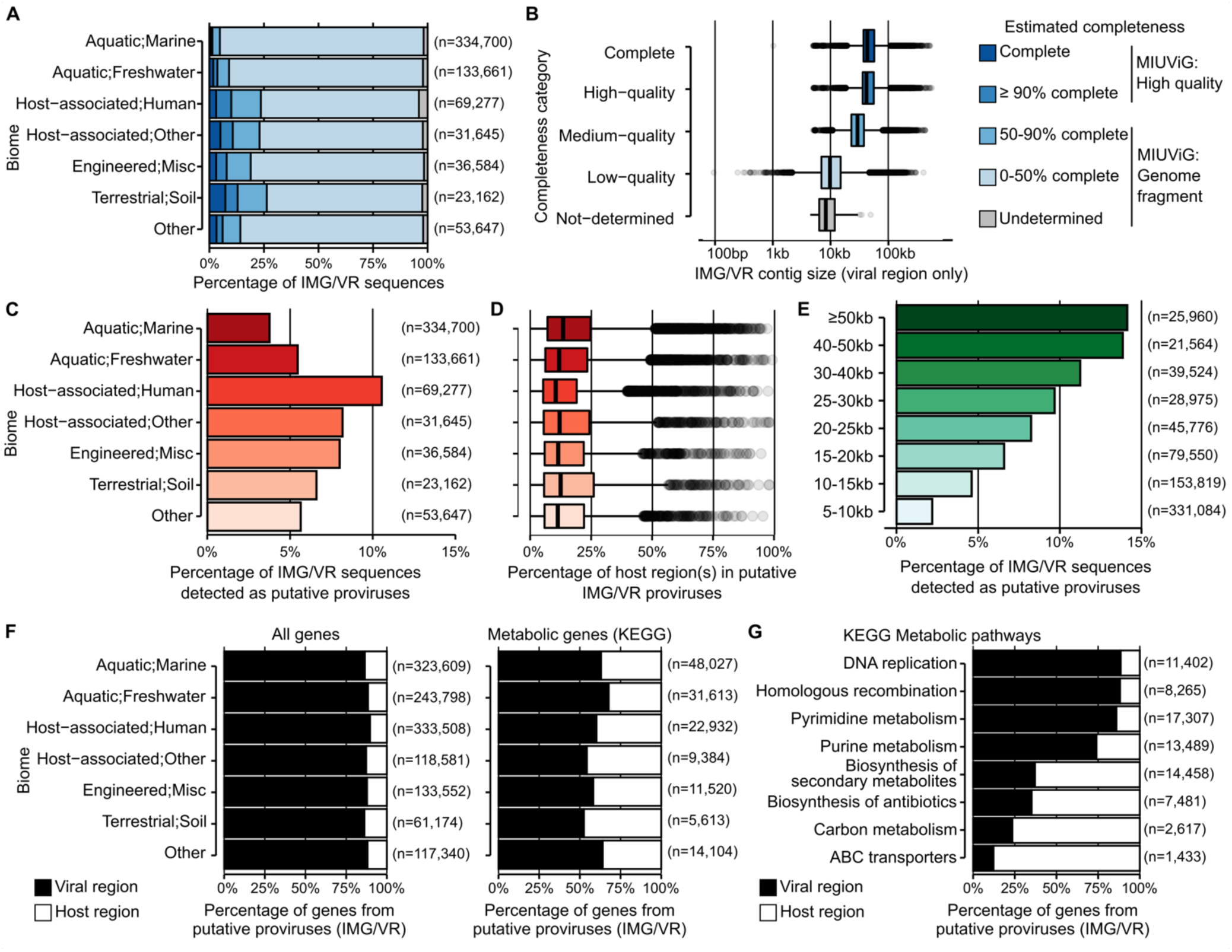
Application of CheckV to the IMG/VR database. A) Estimated completeness of IMG/VR contigs by biome. B) Distribution of IMG/VR contig length across quality tiers. For proviruses, only the size of the predicted viral region was considered. C) Proportion of IMG/VR contigs predicted as proviruses by biome. D) Length of the predicted host region by biome for IMG/VR contigs predicted as proviruses. The region length is indicated as a percentage of the total contig length. E) Proportion of contigs predicted as proviruses by contig length. F) Percentage of all genes from predicted proviruses found in viral/host regions. G) Percentage of metabolic genes from predicted proviruses found in viral/host regions. H) Percentage of genes from selected KEGG pathways for predicted proviruses found in viral/host regions.

### Using CheckV to discriminate viral-encoded functions from host contamination

Finally, we used CheckV to identify putative proviruses from the IMG/VR database that were flanked on one or both sides by host genes. Overall, only 5.2% of contigs followed this pattern (Figure 4C) with 97.1% of host regions occurring on only one side and typically representing a minor fraction of the contig’s length (median=12.0%, Figure 4D). Proviruses were detected in all biomes, although more frequently in host-associated metagenomes. However, longer contigs were more likely to contain a host region (Figure 4D), which may be explained by higher sensitivity of CheckV for longer sequences (Figure S4) and a greater chance of intersecting a host-provirus boundary. Supporting these predictions, the majority of long proviruses (>50 kb with >20% contamination) were confirmed by either VirSorter or VIBRANT (617/805, 76.6%) and often contained integrases (686/805, 85.2%). We also used CheckV to identify proviruses in the GOV dataset, revealing similar patterns (Figure S7). Together, these results confirm that the majority of IMG/VR and GOV sequences are entirely viral or encode a short host-derived region.

Notably, even a small amount of contamination by host-derived sequences can impair downstream analyses, especially ones related to the gene content and functional potential of uncultivated viruses [16]. To illustrate this potential issue, we functionally annotated IMG/VR proviruses using the KEGG database [41] and compared the functions of genes in host versus viral regions. Overall, host regions represented only 12.0% of the genes, but 42.3% of genes assigned to a KEGG metabolic pathway (Figure 4F). Many pathways were highly enriched in host genes, including those for biosynthesis of antibiotics and ABC transporters (Figure 4G and Table S12). For example, 625 genes from the IMG/VR database were annotated as multi-drug resistance efflux pumps, but 92.8% of these were found in host regions. In contrast, KEGG pathways for recombination, mismatch repair, and nucleotide biosynthesis were all enriched in viral regions. Without the detection of provirus boundaries provided by CheckV, it would not have been possible to discriminate truly viral-encoded functions from host contamination except through manual curation, which at the scale of IMG/VR’s data size becomes virtually impossible.

## Discussion

Here we presented CheckV, an automated pipeline for assessing the quality of metagenome-assembled viral genomes along with an expanded database of complete viral genomes we systematically identified from environmental data sources. We anticipate CheckV will be broadly useful in future viral metagenomics studies and for reporting quality statistics required in the MIUViG checklist [17]. Estimation of completeness will be especially valuable to distinguish near-complete genomes from short genome fragments, as these two types of sequences are associated with different limitations and biases. Meanwhile, the removal of genes originating from the host genome will be critically important for reducing false positives in viral studies focusing on auxiliary metabolic genes or novel protein families. We also expect that CheckV’s database of complete viral genomes will be a useful community resource that contains a wealth of untapped insights about novel viruses from diverse environments.

While this first version of CheckV represents a major advance, several improvements may be possible in the future. First, it will be important to incorporate new viral genomes as they become available in order to continually expand the environmental and taxonomic diversity of the reference database. Second, metagenomic read mapping could be used to identify circular contigs, refine virus-host boundaries, and determine genome termini. Third, viral bins and segmented viral genomes, which are represented by multiple sequences, pose several additional challenges not addressed by the current version of CheckV. Finally, we largely focused on bacterial and archaeal viruses in our benchmarking and applications, so modifications may be necessary to extend CheckV for eukaryotic viruses, particularly those with segmented genomes or those that contain many metabolic genes.

## Methods

### Database of HMMs for classification of viral and microbial genes

We selected HMMs from existing databases that could be leveraged to classify genes as either viral or microbial with high specificity. First, 125,754 HMMs were downloaded four databases: VOGDB (release 97, N=25,399, http://vogdb.org), IMG/VR (downloaded January 2020, N=25,281) [37], RVDB (release 17, N=9,911) [42], KEGG Orthology (October 02, 2019 release, N=22,746) [41], Pfam A (release 32, N=17,929) [43], Pfam B (release 27, N=20,000) [44], and TIGRFAM (release 15, N=4,488) [45]. Next, we used hmmsearch v3.1b2 [46] to align the HMMs versus 1,590,764 proteins from 30,903 NCBI GenBank viral genomes (downloaded June 1, 2019) [28] and 5,749,148 proteins from 2,015 bacterial and 239 archaeal genomes from the Genome Taxonomy Database (GTDB; Release 89) [36]. For GTDB, one genome was selected per family and when multiple genomes were available, we chose the one with the highest CheckM quality score (completeness - 5 x contamination). Additionally, we ran VIBRANT v1.2.0 [11], VirSorter v.1.0.5 [10], and PhiSpy v.3.7.8 [20] using default parameters to identify and remove 590,484 viral proteins identified on proviruses in the selected GTDB genomes.

Based on the hmmsearch results, we calculated the percentage of viral and microbial genes matching each HMM at bit-score cutoffs ranging from 25 to 1,000 in increments of 5. We then selected the lowest bit-score cutoff for each HMM that resulted in a >100-fold difference between the percentage of the total viral gene set and the percentage of the total microbial gene set matched by the HMM (i.e. bit-score cutoff for which the hits were either strongly enriched in viruses or microbial genes). To limit false positives, we excluded HMMs that were classified as microbial-specific but were derived from primarily viral databases (VOGDB, IMG/VR, RVDB) or contained viral terms (viral, virus, virion, provirus, capsid, terminase) for HMMs from other databases. Using this approach, 114,765 HMMs were identified as viral-specific or microbial-specific.

Next, we selected the maximally informative subset of HMMs to reduce the size of the database and limit CheckV computing time. First, we retained 44,415 HMMs with at least 20 viral hits or with at least 100 microbial hits after applying the bit-score cutoffs. Next, we calculated the Jaccard similarity between all pairs of HMMs based on each HMMs set of gene hits. For computational efficiency, we used the ‘all_pairs’ function in the SetSimilaritySearch Python package (https://github.com/ekzhu/SetSimilaritySearch). Jaccard similarities were used as input for single-linkage clustering with a Jaccard similarity cutoff of 0.5, resulting in 15,958 non-redundant HMMs (8,773 viral-specific, 7,185 microbial-specific). To form the final database, we selected the HMM with the greatest number of gene hits from each cluster of HMMs.

### Identification of virus-host boundaries

Given a viral contig, CheckV predicts host-virus boundaries in three stages. First, proteins are predicted using Prodigal v2.6.3 (option ‘-p’ for metagenome mode) [47] and compared to the 15,958 HMMs using hmmsearch. Each protein is classified as viral, microbial, or unannotated according to its top-scoring hit after applying the HMM-specific bit-score cutoffs. Additionally, the GC content of each gene is calculated. Second, CheckV evaluates each intergenic region as a potential boundary between a host and a viral region as follows. For each intergenic region, two windows of up to 40 genes are compared, one upstream and one downstream of the potential breakpoint. Windows can contain fewer than 40 genes only if they start or end at a contig boundary. For each window, viral-annotated genes are assigned a score of +1, microbial-annotated genes are assigned a score of −1, and a mean viral score, *V*, across the window is calculated, ignoring unannotated genes. Then, CheckV computes a breakpoint score, *S*, based on the absolute difference in the mean viral score, *V*, and average GC content, *G*, between upstream (5’) and downstream (3’) windows: *S* = |*V*_5′_ − *V*_3′_ | + 0.02 * |*G*_5′_ − *G*_3′_ |. The value of *S* ranges from 0 to 4, given that | *V*_5′_ − *V*_3′_ | and 0.02 * |*G*_5′_ − *G*_3′_ | both range from 0.0 to 2.0. CheckV also stores the orientation of each breakpoint (i.e. host-virus or virus-host) based on the values of *V*_5′_ and *V*_3′_. These scores are computed at each intergenic position, moving from the 5’ end to the 3’ end of a contig. CheckV then identifies significant breakpoints having scores ≥ 1.2, ≥ 30% of genes annotated as microbial in either window, and a total of 4 annotated genes between windows. After these filters, CheckV groups together adjacent breakpoints with the same orientation and, for each group, chooses the breakpoint with the highest score. The algorithm parameters were fine-tuned empirically based on a dataset of mock proviruses and sequences from the IMG/VR database.

### AAI-based estimation of genome completeness

Given a viral contig, CheckV estimates genome completeness in four stages. First, CheckV performs an amino acid alignment of Prodigal-predicted protein coding genes from the contig against the database of reference genomes using DIAMOND [48] with options ‘--evalue 1e-5 -- query-cover 50 --subject-cover 50 -k 10000’. Based on these alignments, the following metrics are computed for the viral contig versus each reference genome: average amino acid identity (AAI): length-weighted average identity across aligned proteins, alignment fraction (AF): the percent of amino acids aligned from the query sequence, and alignment score: AAI multiplied by AF. Second, CheckV identifies the top hit in the database for the contig (i.e. reference genome with the highest alignment score) and all reference genomes with alignment scores that are within 50% of the top hit. The expected genome length of the viral contig, *Ĝ*, is then estimated by taking a weighted average of the genome sizes of matched reference genomes, where the alignment scores are used as weights. Reference genome lengths are further weighted based on their source: 2.0 for isolate viruses and 1.0 for metagenome-derived viruses, which are more likely to contain assembly errors and artifacts. CheckV also reports the confidence level of this estimate (low, medium, high), which is determined based on the length of the viral contig and the alignment score to the top reference genome. (See below for more details on how confidence levels are estimated). Third, CheckV estimates the genome completeness of each viral contig, *Ĉ*, using the formula: *Ĉ* = 100 * *L*/ *Ĝ*, where *L* is the length of the viral region for proviruses, or the contig length otherwise.

### HMM-based estimation of genome completeness

An HMM-based approach was developed to estimate completeness for novel viruses that are too diverged from CheckV genomes to obtain an AAI-based estimate. First, CheckV identifies viral genes on the contig based on comparison to the 8,773 viral HMMs (as described in ‘Identification of virus-host boundaries’). Each HMM is associated with one or more reference genome lengths (based on CheckV references where the HMM is found) and the coefficient of variation (CV) is computed, in order to prioritize HMMs with low genome length variability. For each contig, CheckV identifies the viral HMM with the lowest CV that is associated with at least 10 reference genomes. Next, the program compares the contig length to the distribution of genome lengths represented by the selected HMM and chooses the level of completeness (0-100%) that would place the contig in the upper 5th percentile of the genome length distribution. Thus, the approach is designed to estimate the maximum level of genome completeness that can be ascertained with at least 95% probability.

### Confidence levels for AAI-based completeness estimates

We conducted a large-scale benchmarking experiment to derive confidence levels for AAI-based completeness estimation. First, we extracted a random fragment from each of CheckV’s reference genomes to simulate metagenomic contigs of varying length (200 bp, 500 bp, 1 kb, 2 kb, 5 kb, 10 kb, 20 kb, and 50 kb). Next, we used CheckV to compute the alignment score between each contig and each complete genome in the reference database. We then compared the true genome length of each contig (i.e. the length before fragmentation), *L*, to the estimated genome length based on each matched reference genome, 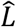, and computed the relative unsigned error, as 100 * 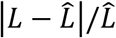. We then computed the median relative unsigned error after grouping the estimates based on their alignment score and contig length. Finally, we determined three confidence levels: high confidence (0-5% median unsigned error, 76.6% mean AAI, 65.0% mean AF); medium confidence (5-10% median unsigned error, 51.8% mean AAI%, 54.7% mean AF); low confidence (>10% median unsigned error, 47.5% mean AAI, 36.6% mean AF. Using this information, CheckV reports a confidence level in the estimated completeness value for each input contig based on the contig length and alignment score (i.e. combination of AAI and AF) to the top database hit. By default, only medium- and high-confidence estimates are included in the final report.

### Database of complete viral genomes for AAI-based completeness estimation

We downloaded 30,903 genomes from NCBI GenBank on June 1, 2019. 1,937 genomes were excluded that were indicated as ‘partial’, ‘chimeric’, or ‘contaminated’. Of the 28,966 that remained, 677 (2.3%) were labeled as ‘metagenomic’ or ‘environmental’, indicating the vast majority are derived from cultivated isolates.

Next, we used CheckV to systematically search for complete genomes of uncultivated viruses from publicly available and previously assembled metagenomes, metatranscriptomes, and metaviromes. An assembled contig was considered complete if it was at least 2,000 bp long and included a direct terminal repeat (DTR) of at least 20 bp (DTR contigs). We searched for DTR contigs in the following datasets: 19,483 metagenomes and metatranscriptomes from IMG/M (accessed on September 2019) [29], 11,752 metagenomes from MGnify (accessed on April 16, 2019) [30], 9,428 metagenomes assembled by Pasolli et al. [32], an expanded collection of 4,763 metagenomes from the HGM dataset [31], 1,831 viromes from the HuVirDB [33], and 145 viromes from the global ocean [6].

From this initial search, we identified a total of 751,567 DTR contigs. To minimize false positives and other artifacts, we removed the following: (1) 45,448 contigs with low complexity repeats (e.g. AAAAA…), as determined by dustmasker from the BLAST+ package v2.9.0 [49], (2) 11,359 contigs classified as a provirus by CheckV (e.g. flanked by host region), (3) 5,737 contigs with repetitive repeats occurring more than 5 times per contig, which could represent repetitive genetic elements like CRISPR arrays, (4) 6,543 contigs that contained a large duplicated region spanning >=20% of the contig length which results from the rare instances where assemblers concatenate multiple copies of the same genome, and (5) 1,293 contigs containing >=1% ambiguous base calls. After these filters, 686,030 contigs remained (91.3% of total).

Next, we used a combination of CheckV marker genes and VirFinder [9] to classify 116,666 DTR contigs as viral. First, the DTR contigs were used as input to VirFinder v1.1 with default parameters and to CheckV to identify viral and microbial marker genes. We additionally searched for genes related to plasmids and other non-viral mobile genetic elements using a database of 141 HMMs from recent publications [50-52]. A contig was classified as viral if the number of viral genes exceeded number of microbial and plasmid genes (N=99,345), or VirFinder reported a p-value < 0.01 with no plasmid genes and <= 1 microbial gene (N=36,084).

### Taxonomic annotation of CheckV reference genomes

Annotations were determined based on HMM searches against a custom database of 1,000 taxonomically informative HMMs from the VOG database. These HMMs were selected for major bacterial and archaeal viral groups with consistent genome length and at least 10 representative genomes, including: *Caudovirales*, CRESS-DNA and *Parvoviridae, Autolykiviridae, Fusello-* and *Guttaviridae, Inoviridae, Ligamenvirales Ampulla-Bicauda-* and *Turriviridae, Microviridae*, and *Riboviria*. For each group, VOGs found in ≥10% of the group members and never detected outside of this group were considered as marker genes. All CheckV reference genomes were annotated based on the clade with the most HMM hits. Overall, 96.4% of HMM hits were to a single viral taxon.

### Validating the completeness of CheckV reference genomes

Next, we validated the completeness for all GenBank genomes and DTR contigs. First, we used CheckV to estimate the completeness for all sequences after excluding self matches. This was performed using a database of just GenBank sequences and another of just DTR contigs. Any sequence with <90% estimated completeness using either database was excluded (medium- and high-confidence estimates only). Second, we compared the genome length to the known distribution of genome lengths for the annotated viral taxon (e.g. *Microviridae*). Any genome considered an outlier or shorter than the shortest reference genome for the annotated clade was excluded. After these filters, we then selected genomes with ≥ 90% estimated completeness using either database (medium- and high-confidence estimates only) or longer than 30 kb without a completeness estimate. These selection criteria were chosen to minimize the number of false positives (i.e. genome fragments wrongly considered complete genomes) at the cost of some false negatives (i.e. removing truly complete genomes). This resulted in 24,834 GenBank genomes and 76,262 DTR contigs that were used to form the final CheckV genome database.

### Generating a non-redundant set of CheckV reference genomes

Average nucleotide identity (ANI) and the alignment fraction (AF) were computed between the 24,834 GenBank genomes and 76,262 DTR contigs using a custom script. Specifically, we used blastn from the blast+ package v.2.9.0 (options: perc_identity=90 max_target_seqs=10000) to generate local alignments between all pairs of genomes. Based on this, we estimated ANI as the average DNA identity across alignments after weighting the alignments by their length. The alignment fraction (AF) was computed by taking the total length of merged alignment coordinates and dividing by the length of each genome. Clustering was then performed using a greedy, centroid-based algorithm in which (1) genomes were sorted by length, (2) the longest genome was designated as the centroid of a new cluster, (3) all genomes within 95% ANI and 85% AF were assigned to that cluster, and steps (2-3) were repeated until all genomes had been assigned to a cluster, resulting in 52,141 non-redundant genomes.

### Benchmarking CheckV and comparison to existing tools

To benchmark CheckV’s detection of host regions, we constructed a mock dataset of proviruses. 382 viral genomes were downloaded from NCBI GenBank (after June 1, 2019) and paired with 76 GTDB genomes (71 from bacterial 5 from archaea). The pairing was performed at the genus level based on the annotated name of the virus and host (e.g. *Escherichia* phage paired with *Escherichia* bacterial genome). When multiple GTDB genomes were available for a given bacterial genus, we selected the one with the highest CheckM quality score, and we selected a maximum of 10 GenBank genomes per bacterial genus to reduce the influence of a few overrepresented groups. Any GenBank or GTDB genome that was used at any stage for training CheckV was excluded. Proviruses were simulated at varying contig lengths (5, 10, 20, 50, 100 kb) with varying levels of host contamination (10, 20, 50%; defined as the % of the contig length derived from the microbial genome). Microbial genome fragments were appended to either the 5’ or 3’ end of the viral fragment at random. As a negative control, we also simulated contigs that were entirely viral (i.e. no flanking microbial region) at the same contig lengths.

Mock proviruses were used as input to CheckV using default parameters. For comparison, we also ran VIBRANT v1.2.0 [11], VirSorter v.1.0.5 [10], PhiSpy v.3.7.8 [20], and Phigaro v0.1.5.0 [21]. All tools were run with default options, with the exception of VIBRANT and VirSorter, which were run with the ‘--virome’ flag to increase sensitivity. Nucleotide sequences were used as input to all tools, except Phispy, for which we first ran Prokka v1.14.5 [53] to generate the required input file. A contig was classified as a provirus if it contained a predicted viral region that covered <95% of its length. Each prediction was then classified as a true positive (TP; provirus classified as provirus), false positive (FP; viral contig classified as provirus), true negative (TN; viral contig not classified as provirus), and false negative (FN; provirus classified as provirus). For the TPs, we also compared the true length of the host region to the predicted length.

We used the same 382 provirus genomes to benchmark CheckV’s completeness estimation. For each of the 382 genomes, we extracted 10 contigs at a random start position and with a completeness value between 1% and 100%. The completeness of all contigs was estimated using CheckV with default parameters. VIBRANT v1.2.0 was also run using default parameters to assign contigs to quality tiers.

### Application of CheckV to the IMG/VR and GOV datasets

We downloaded 735,106 contigs longer than 5 kb from IMG/VR 2.0 [37], after excluding viral genomes from cultivated isolates and proviruses identified from microbial genomes. We also downloaded 488,131 contigs longer than 5 kb or circular from the Global Ocean Virome (GOV) 2.0 dataset [6] (datacommons.cyverse.org/browse/iplant/home/shared/iVirus/GOV2.0). These were used as input to CheckV to estimate the completeness, identify host-virus boundaries, and predict closed genomes. When running the completeness module, we excluded perfect matches (100% AAI and 100% AF) to prevent any DTR contig from matching itself in the database (since IMG/VR 2.0 and GOV 2.0 were used as data sources to form the CheckV database). A Circos plot [54] was used to link IMG/VR contigs to their top matches in CheckV database. Protein coding genes were predicted from proviruses using Prodigal and compared to HMMs from KEGG Orthology (October 02, 2019 release) using hmmsearch from the HMMER package v3.1b2 (e-value ≤ 1e-5 and score ≥ 30). Pfam domains with the keyword “integrase” and “recombinase” were also identified across all proviruses.

The largest DTR contig we identified from IMG/VR and included in the CheckV database was further annotated to illustrate the type of virus and genome organization represented (IMG ID = 3300025697_Ga0208769_1000001). CDS prediction and functional annotations were obtained from IMG [29]. Annotation for provirus hallmark genes including a terminase large subunit (TerL) and major capsid protein were confirmed via HHPred [55] (databases included PDB 70_8, SCOPe70 2.07, Pfam-A 32.0, and CDD 3.18, score > 98). A circular genome map was drawn with CGView [56]. To place this contig in an evolutionary context, we built a TerL phylogeny including the most closely related sequences from a global search for large phages [13]. The TerL amino acid sequence from the DTR contig was compared to all TerL sequences from the “huge proviruses” dataset via blastp (e-value ≤ 1e-05 and score ≥ 50) to identify the 30 most similar sequences (sorted based on blastp bit-score). These reference sequences and 3300025697_Ga0208769_1000001 were aligned with MAFFT v7.407 [57] using default parameters, the alignment automatically cleaned with trimAL v1.4.rev15 with the --gappyout option [58], and a phylogeny built with IQ-Tree v1.5.5 with built-in model selection (optimal model suggested: LG+R4) [59]. The resulting tree was visualized with iToL [60].

## Data and code availability

CheckV is written in Python and is freely available as open source software at https://bitbucket.org/berkeleylab/CheckV under a BSD license. CheckV quality statistics and listing of complete viral genomes from IMG/VR will be available in the next version scheduled for release on September 2020.

## Contributions

S.N. and S.R conceived of the project. S.N. and S.R. performed experiments and analyzed data. S.N. built databases and designed algorithms. S.N. and A.P.C. wrote the code. A.P.C. packaged the software. S.N. and S.R. drafted the manuscript with feedback from A.P.C, E.E.F., and N.K. N.K. supervised the project. All authors reviewed and approved the manuscript.

## Acknowledgements

We thank Professor Riccardo Cavicchioli for providing additional information on the sample from which the largest DTR contig was assembled. The work conducted by the U.S. Department of Energy Joint Genome Institute is supported by the Office of Science of the U.S. Department of Energy under contract no. DE-AC02-05CH11231. This work was also supported the grant #2016/23218-0 from São Paulo Research Foundation (FAPESP). A.P.C. received a scholarship #2018/04240-0 from FAPESP.

## References

1. Koonin, E.V., et al., Global Organization and Proposed Megataxonomy of the Virus World. Microbiol Mol Biol Rev, 2020. 84(2).

2. Shkoporov, A.N. and C. Hill, Bacteriophages of the Human Gut: The “Known Unknown” of the Microbiome. Cell Host Microbe, 2019. 25(2): p. 195–209.

3. Williamson, K.E., et al., Viruses in Soil Ecosystems: An Unknown Quantity Within an Unexplored Territory. Annu Rev Virol, 2017. 4(1): p. 201–219.

4. Breitbart, M., et al., Phage puppet masters of the marine microbial realm. Nat Microbiol, 2018. 3(7): p. 754–766.

5. Paez-Espino, D., et al., Uncovering Earth’s virome. Nature, 2016. 536(7617): p. 425–30.

6. Gregory, A.C., et al., Marine DNA Viral Macro- and Microdiversity from Pole to Pole. Cell, 2019. 177(5): p. 1109–1123 e14.

7. Gregory, A.C., et al., The human gut virome database. bioRxiv, 2019.

8. Emerson, J.B., et al., Host-linked soil viral ecology along a permafrost thaw gradient. Nat Microbiol, 2018. 3(8): p. 870–880.

9. Ren, J., et al., VirFinder: a novel k-mer based tool for identifying viral sequences from assembled metagenomic data. Microbiome, 2017. 5(1): p. 69.

10. Roux, S., et al., VirSorter: mining viral signal from microbial genomic data. PeerJ, 2015. 3: p. e985.

11. Kieft, K., Z. Zhou, and K. Anantharaman, VIBRANT: Automated recovery, annotation and curation of microbial viruses, and evaluation of virome function from genomic sequences. bioRxiv, 2019.

12. Philippe, N., et al., Pandoraviruses: amoeba viruses with genomes up to 2.5 Mb reaching that of parasitic eukaryotes. Science, 2013. 341(6143): p. 281–6.

13. Al-Shayeb, B., et al., Clades of huge phages from across Earth’s ecosystems. Nature, 2020. 578(7795): p. 425–431.

14. Smits, S.L., et al., Assembly of viral genomes from metagenomes. Front Microbiol, 2014. 5: p. 714.

15. Roux, S., et al., Minimum Information about an Uncultivated Virus Genome (MIUViG). Nat Biotechnol, 2019. 37(1): p. 29–37.

16. Roux, S., et al., Assessment of viral community functional potential from viral metagenomes may be hampered by contamination with cellular sequences. Open Biol, 2013. 3(12): p. 130160.

17. Parks, D.H., et al., CheckM: assessing the quality of microbial genomes recovered from isolates, single cells, and metagenomes. Genome Res, 2015. 25(7): p. 1043–55.

18. Belyi, V.A., A.J. Levine, and A.M. Skalka, Sequences from ancestral single-stranded DNA viruses in vertebrate genomes: the parvoviridae and circoviridae are more than 40 to 50 million years old. J Virol, 2010. 84(23): p. 12458–62.

19. Chung, C.H., et al., Predicting genome terminus sequences of Bacillus cereus-group bacteriophage using next generation sequencing data. BMC Genomics, 2017. 18(1): p. 350.

20. Akhter, S., R.K. Aziz, and R.A. Edwards, PhiSpy: a novel algorithm for finding prophages in bacterial genomes that combines similarity- and composition-based strategies. Nucleic Acids Res, 2012. 40(16): p. e126.

21. Starikova, E.V., et al., Phigaro: high throughput prophage sequence annotation. 2019.

22. Hindmarsh, P. and J. Leis, Retroviral DNA integration. Microbiol Mol Biol Rev, 1999. 63(4): p. 836–43, table of contents.

23. Tisza, M.J., et al., Discovery of several thousand highly diverse circular DNA viruses. Elife, 2020. 9.

24. Casjens, S.R. and E.B. Gilcrease, Determining DNA packaging strategy by analysis of the termini of the chromosomes in tailed-bacteriophage virions. Methods Mol Biol, 2009. 502: p. 91–111.

25. Munoz-Lopez, M. and J.L. Garcia-Perez, DNA transposons: nature and applications in genomics. Curr Genomics, 2010. 11(2): p. 115–28.

26. Yan, Z., et al., Inverted terminal repeat sequences are important for intermolecular recombination and circularization of adeno-associated virus genomes. J Virol, 2005. 79(1): p. 364–79.

27. Savilahti, H. and D.H. Bamford, Linear DNA replication: inverted terminal repeats of five closely related Escherichia coli bacteriophages. Gene, 1986. 49(2): p. 199–205.

28. Sayers, E.W., et al., GenBank. Nucleic Acids Res, 2020. 48(D1): p. D84–D86.

29. Chen, I.A., et al., IMG/M v.5.0: an integrated data management and comparative analysis system for microbial genomes and microbiomes. Nucleic Acids Res, 2019. 47(D1): p. D666–D677.

30. Mitchell, A.L., et al., MGnify: the microbiome analysis resource in 2020. Nucleic Acids Res, 2020. 48(D1): p. D570–D578.

31. Nayfach, S., et al., New insights from uncultivated genomes of the global human gut microbiome. Nature, 2019. 568(7753): p. 505–510.

32. Pasolli, E., et al., Extensive Unexplored Human Microbiome Diversity Revealed by Over 150,000 Genomes from Metagenomes Spanning Age, Geography, and Lifestyle. Cell, 2019. 176(3): p. 649–662 e20.

33. Soto-Perez, P., et al., CRISPR-Cas System of a Prevalent Human Gut Bacterium Reveals Hyper-targeting against Phages in a Human Virome Catalog. Cell Host Microbe, 2019. 26(3): p. 325–335 e5.

34. Mukherjee, S., et al., Genomes OnLine database (GOLD) v.7: updates and new features. Nucleic Acids Res, 2019. 47(D1): p. D649–D659.

35. Mauri, M., et al., RAWGraphs, In Proceedings of the 12th Biannual Conference on Italian SIGCHI Chapter - CHItaly ‘17. 2017. p. 1–5.

36. Parks, D.H., et al., A standardized bacterial taxonomy based on genome phylogeny substantially revises the tree of life. Nat Biotechnol, 2018.

37. Paez-Espino, D., et al., IMG/VR v.2.0: an integrated data management and analysis system for cultivated and environmental viral genomes. Nucleic Acids Res, 2019. 47(D1): p. D678–D686.

38. Bobay, L.M., M. Touchon, and E.P. Rocha, Pervasive domestication of defective prophages by bacteria. Proc Natl Acad Sci U S A, 2014. 111(33): p. 12127–32.

39. Rinke, C., et al., Validation of picogram- and femtogram-input DNA libraries for microscale metagenomics. PeerJ, 2016. 4: p. e2486.

40. Warwick-Dugdale, J., et al., Long-read viral metagenomics captures abundant and microdiverse viral populations and their niche-defining genomic islands. PeerJ, 2019. 7: p. e6800.

41. Kanehisa, M. and S. Goto, KEGG: Kyoto Encyclopedia of Genes and Genomes. Nucleic Acids Research, 2000. 28(1): p. 27–30.

42. Goodacre, N., et al., A Reference Viral Database (RVDB) To Enhance Bioinformatics Analysis of High-Throughput Sequencing for Novel Virus Detection. mSphere, 2018. 3(2).

43. El-Gebali, S., et al., The Pfam protein families database in 2019. Nucleic Acids Res, 2019. 47(D1): p. D427–D432.

44. Finn, R.D., et al., The Pfam protein families database. Nucleic Acids Res, 2010. 38(Database issue): p. D211–22.

45. Haft, D.H., et al., TIGRFAMs and Genome Properties in 2013. Nucleic Acids Res, 2013. 41(Database issue): p. D387–95.

46. Eddy, S.R., Accelerated Profile HMM Searches. PLoS Comput Biol, 2011. 7(10): p. e1002195.

47. Hyatt, D., et al., Gene and translation initiation site prediction in metagenomic sequences. Bioinformatics, 2012. 28(17): p. 2223–30.

48. Buchfink, B., C. Xie, and D.H. Huson, Fast and sensitive protein alignment using DIAMOND. Nat Methods, 2015. 12(1): p. 59–60.

49. Camacho, C., et al., BLAST+: architecture and applications. BMC Bioinformatics, 2009. 10: p. 421.

50. Jorgensen, T.S., et al., Hundreds of circular novel plasmids and DNA elements identified in a rat cecum metamobilome. PLoS One, 2014. 9(2): p. e87924.

51. Martini, M.C., et al., Genomics of high molecular weight plasmids isolated from an on-farm biopurification system. Sci Rep, 2016. 6: p. 28284.

52. Jorgensen, T.S., et al., Plasmids, Viruses, And Other Circular Elements In Rat Gut. bioRxiv, 2017.

53. Seemann, T., Prokka: rapid prokaryotic genome annotation. Bioinformatics, 2014. 30(14): p. 2068–9.

54. Krzywinski, M., et al., Circos: an information aesthetic for comparative genomics. Genome Res, 2009. 19(9): p. 1639–45.

55. Zimmermann, L., et al., A Completely Reimplemented MPI Bioinformatics Toolkit with a New HHpred Server at its Core. J Mol Biol, 2018. 430(15): p. 2237–2243.

56. Stothard, P. and D.S. Wishart, Circular genome visualization and exploration using CGView. Bioinformatics, 2005. 21(4): p. 537–9.

57. Katoh, K. and D.M. Standley, MAFFT multiple sequence alignment software version 7: improvements in performance and usability. Mol Biol Evol, 2013. 30(4): p. 772–80.

58. Capella-Gutierrez, S., J.M. Silla-Martinez, and T. Gabaldon, trimAl: a tool for automated alignment trimming in large-scale phylogenetic analyses. Bioinformatics, 2009. 25(15): p. 1972–3.

59. Nguyen, L.T., et al., IQ-TREE: a fast and effective stochastic algorithm for estimating maximum-likelihood phylogenies. Mol Biol Evol, 2015. 32(1): p. 268–74.

60. Letunic, I. and P. Bork, Interactive Tree Of Life (iTOL) v4: recent updates and new developments. Nucleic Acids Res, 2019. 47(W1): p. W256–W259.

