## Supplementary Information for "CheckV: assessing the quality of metagenome-assembled viral genomes"

### Supplementary figures

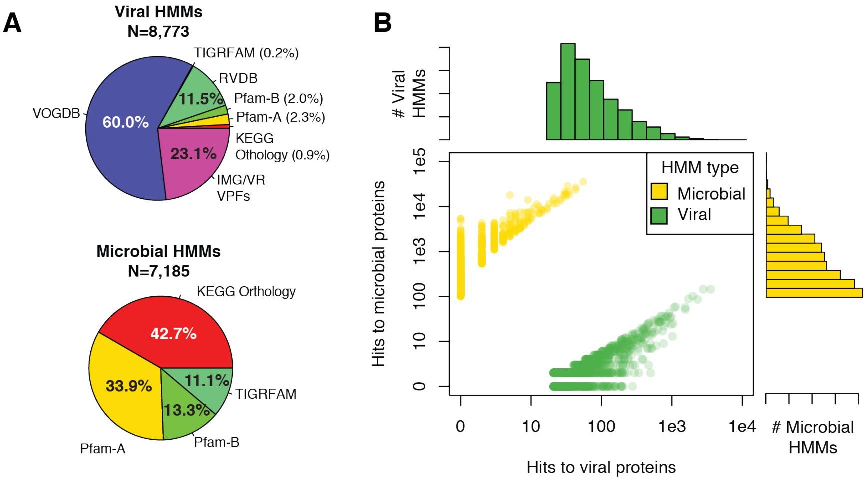


**Figure S1. CheckV's database of viral- and microbial-specific HMMs.** A) Non-redundant viral and microbial HMMs were selected from seven reference databases. B) The distribution of the number of hits to viral and microbial proteins for the CheckV HMMs shown in A.

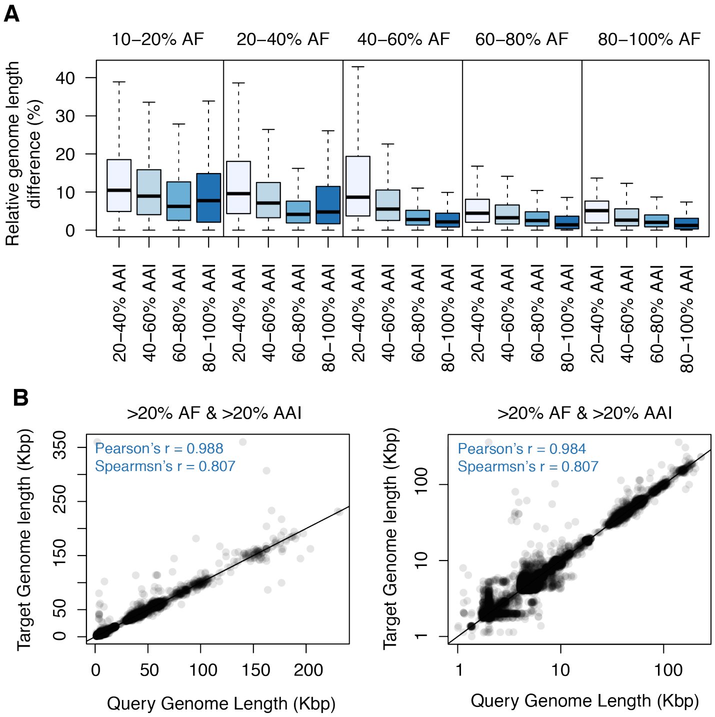


**Figure S2. Variation in genome size between related viruses.** The relatedness between all CheckV reference genomes was estimated based on their average amino acid identity (AAI) and alignment fraction (AF). A) The relative difference in genome length for viruses with varying degrees of relatedness. B) Scatterplots showing genome sizes for related viruses. The right panel shows genome sizes on a log10 scale.

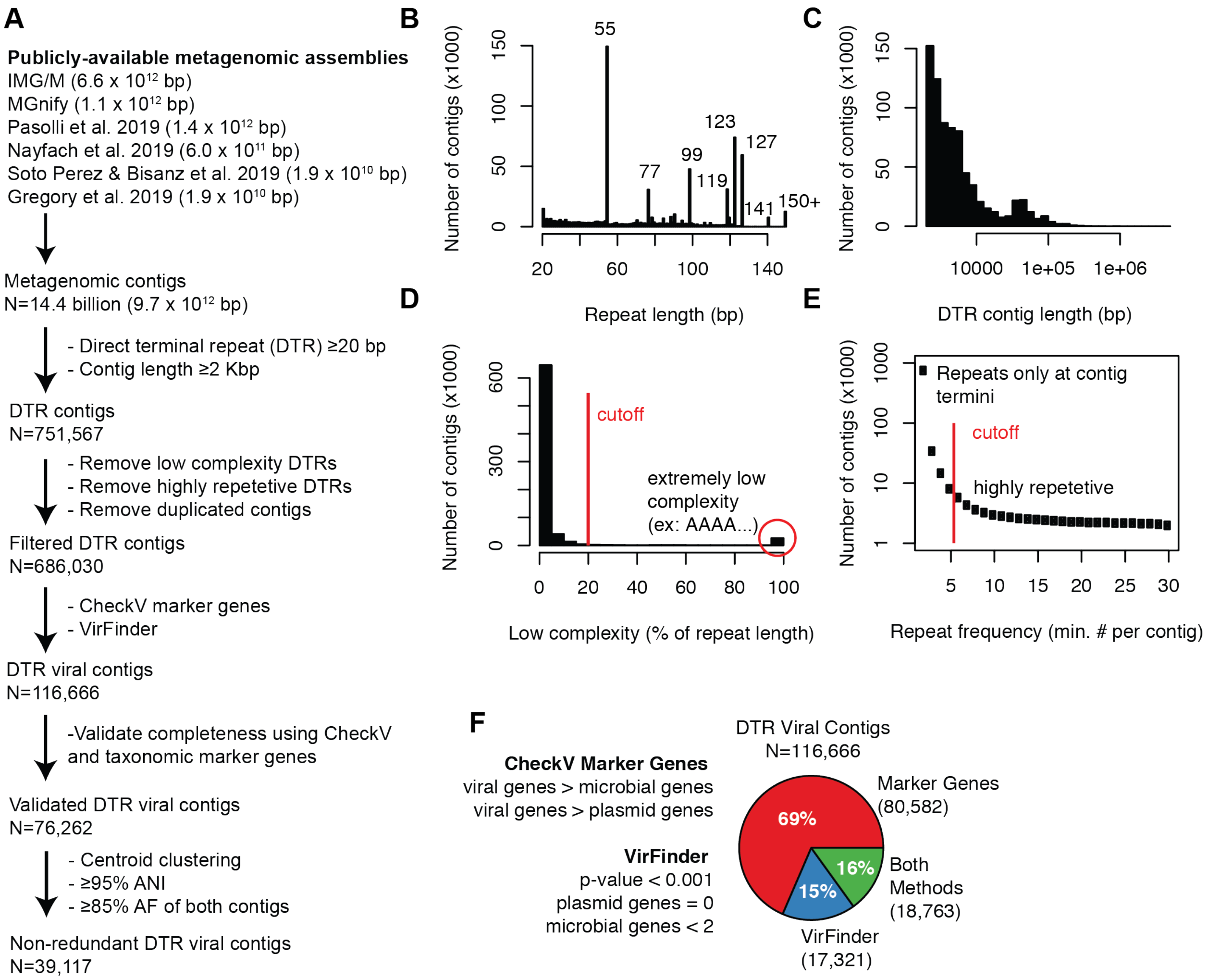


**Figure S3. Identification of viral DTR contigs.** A) Publicly available metagenomes were systematically mined for 76,262 DTR viral contigs, resulting in 39,117 non-redundant contigs after de-replication at 95% ANI over 85% the length of both sequences. B-E) Summary statistics across the 751,567 DTR contigs before filtering. B) Distribution of the length of direct terminal repeats (DTRs). A considerable number of DTRs occur at specific lengths (e.g. 55, 77, 99 bp). These odd-numbered lengths likely correspond with k-mer lengths utilized by various metagenomic assembly tools. When faced with assembling reads from a circular template, they appear to break the contig in a random location and leave behind a repeated sequence at the start and end of the contig equal to the k-mer length. C) The length (log scale) of all DTR contigs. D-E) A small number of contigs are likely false positives due to a low complexity repeat (e.g. AAAAAA…) or a highly repetitive repeat (i.e. occurring not just at termini). F) After removing spurious complete genomes, the DTR contigs were screened for viral signatures, revealing 116,666 viral contigs. These were identified using a combination of CheckV's marker genes, plasmid genes from recent publications, and VirFinder [1].

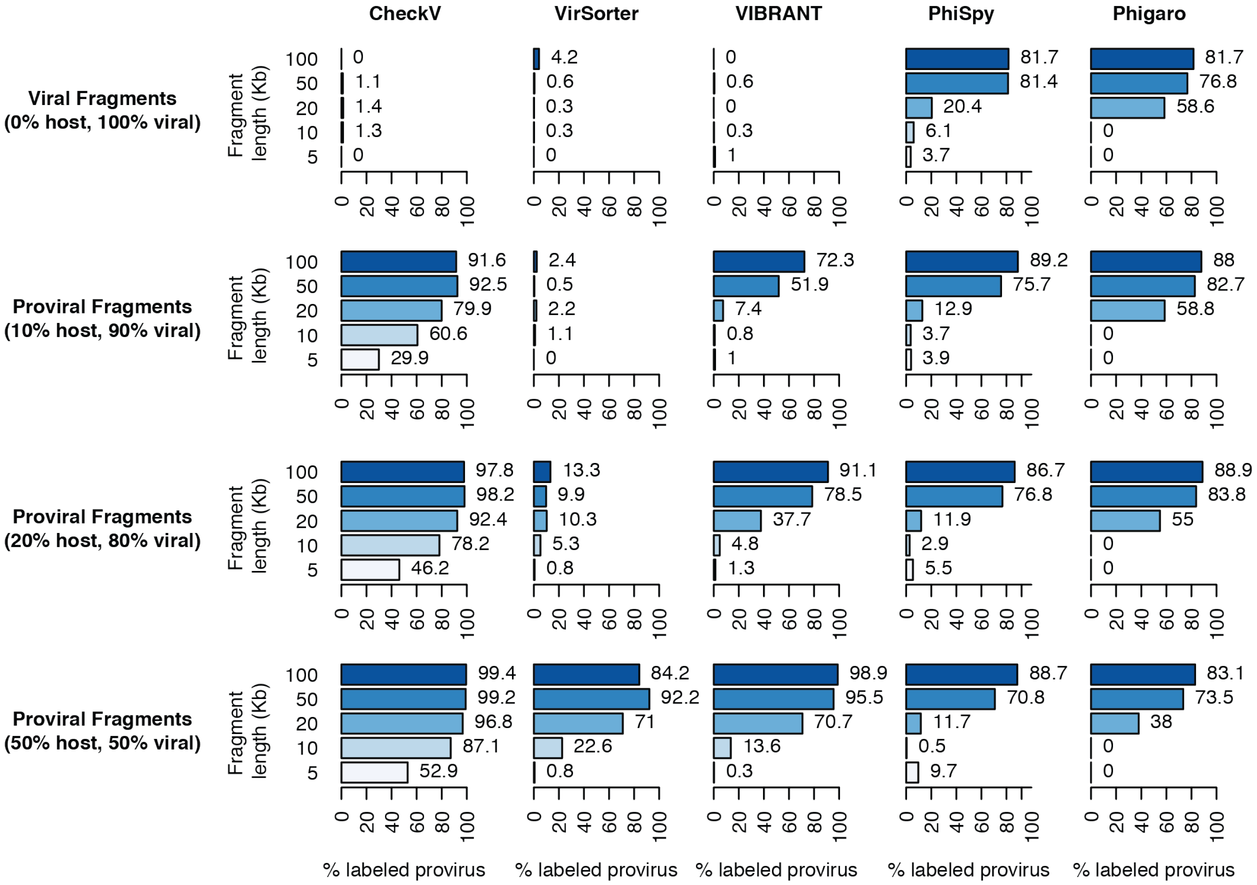


**Figure S4. Provirus classification accuracy for CheckV and other tools.** Proviral genome fragments were generated at various read lengths (5 to 100 kb) and levels of host contamination (0 to 50%) and used as input to CheckV and other tools. A fragment was classified as a provirus if it contained a predicted viral region that covered < 95% of the fragment length.

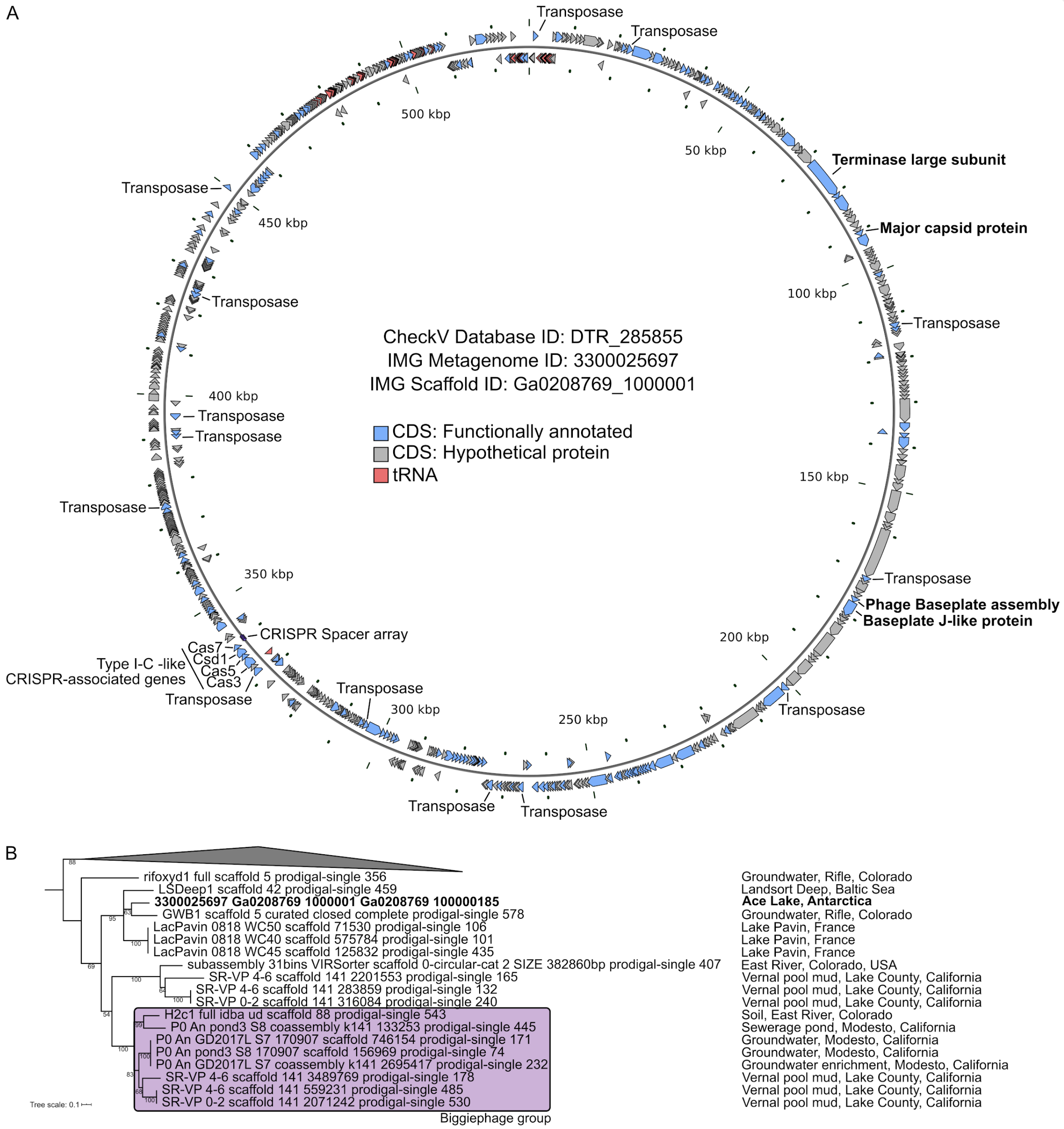


**Figure S5. Genome map and phylogeny of contig Ga0222679_1000001**. A. Genome map of putative circular contig Ga0222679_1000001. Annotations were obtained from IMG [2] and manual annotation of phage proteins (terminase and major capsid protein) via HHPred [3].

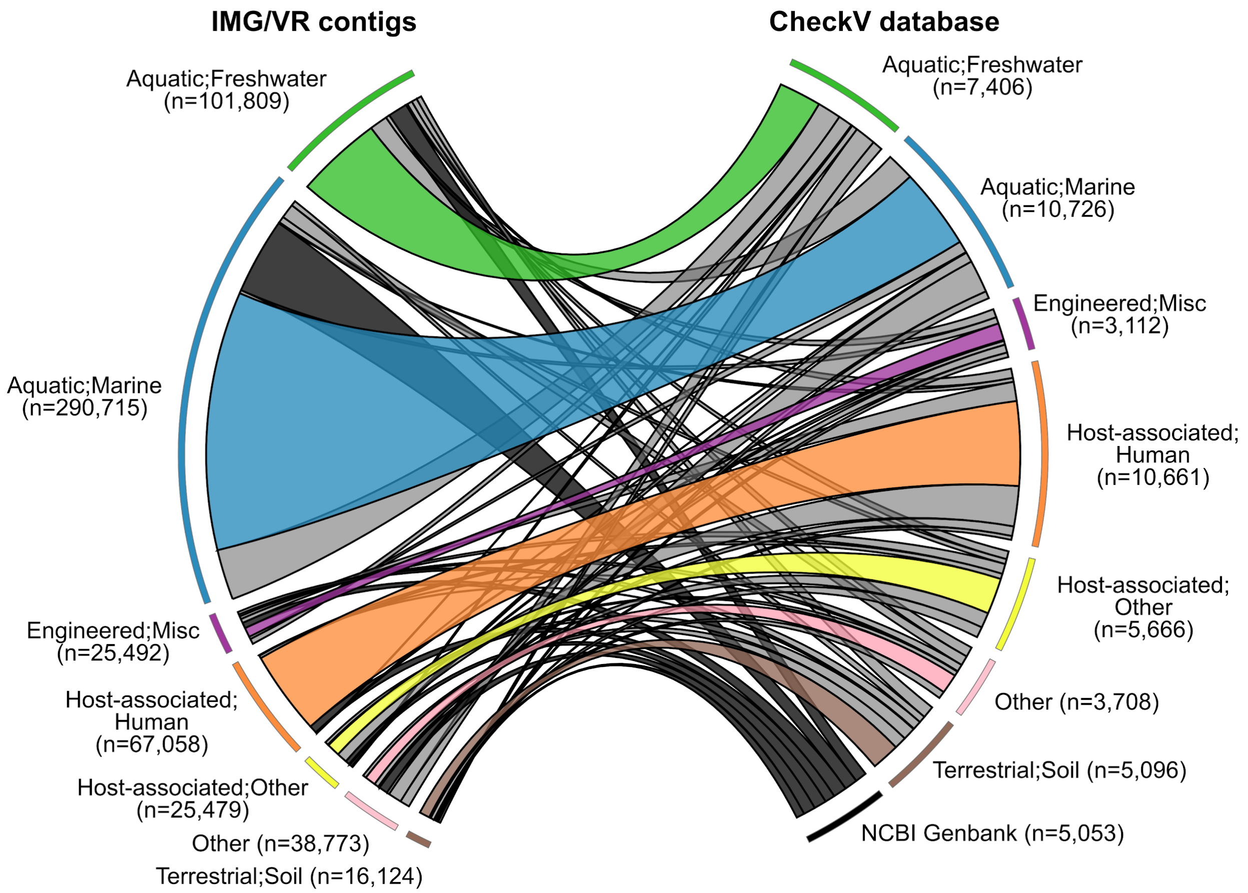


**Figure S6. Association between IMG/VR contigs and CheckV reference genomes.** IMG/VR contigs (left) are classified by the biome of their original metagenomes and connected to the top hit in the CheckV database (right). Cases in which a reference contig is used to estimate the genome of an IMG/VR sequence from the same biome (e.g. marine IMG/VR contig and marine CheckV reference) are colored by biome, while other cases are colored in grey.

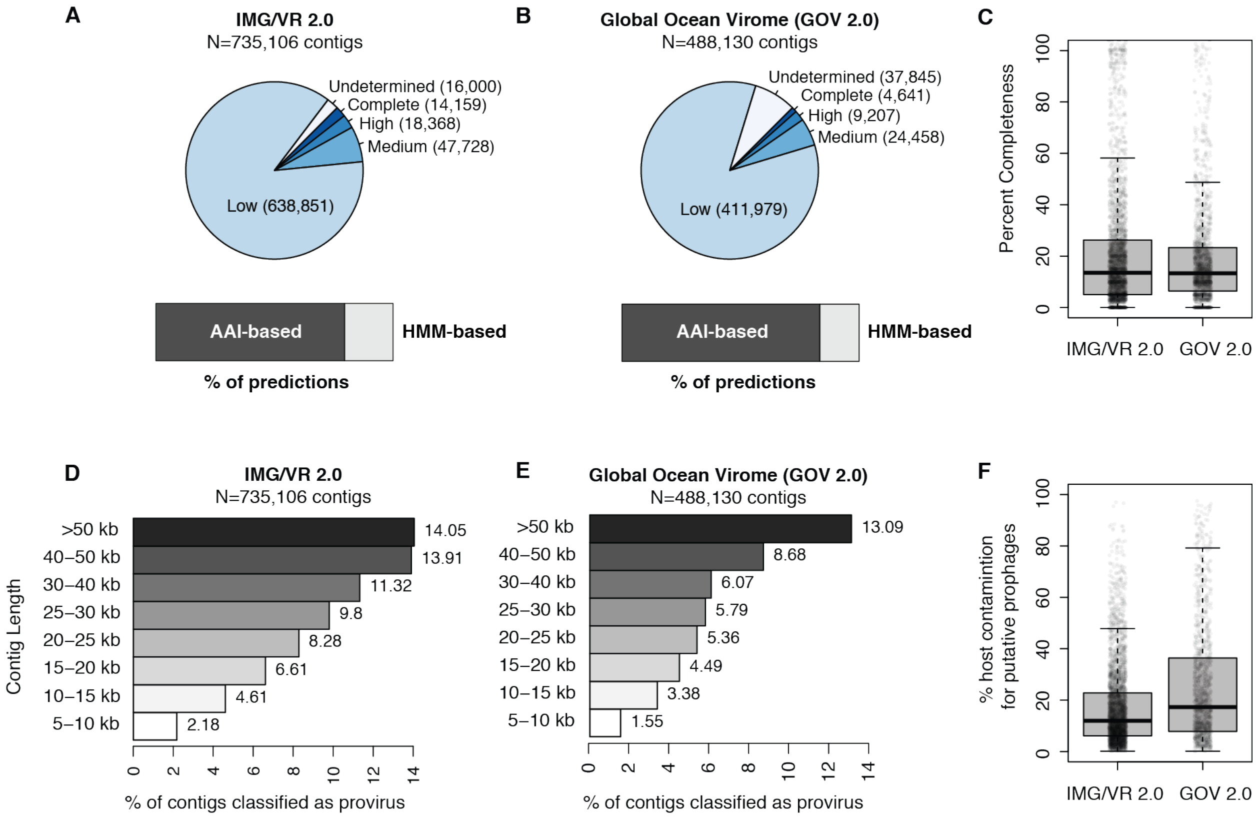


**Figure S7. Application of CheckV to IMG/VR and the Global Ocean Virome datasets.** A) Quality tiers across viral contigs from IMG/VR 2.0 [4] and B) the GOV 2.0 dataset [5]. The bar plots indicate the % of completeness estimates made with the AAI- or HMM-based approaches. C) Distribution of completeness across contigs from each dataset. D) Percent of contigs classified as a provirus for IMG/VR 2.0. and E) for GOV 2.0. F) Host contamination (i.e. percent of length derived from host regions) across datasets.

### Supplementary text

#### Investigating DTR contigs classified as *Retrovirales* and *Riboviria*

Since genomes from *Retrovirales* and *Riboviria* (i.e. RNA viruses) are typically linear, we further analyzed DTR sequences affiliated to these clades to identify putative errors or misannotation. For *Retrovirales*, most sequences with DTR (>97%) were ≤ 15kb, which is consistent with the size range of complete retrovirus genomes. A best blast hit affiliation of these contigs against NCBI Viral RefSeq revealed that the vast majority (>90%) were most similar to *Metaviridae*, i.e. retrotransposon-like with long terminal repeats. The second most common group to which these sequences were affiliated was the *Caulimoviridae* family, with a circular genome. Hence, DTR contigs affiliated to *Retrovirales* seemingly represented genuine complete viral genomes and/or retrotransposons.

For *Riboviria*, >97% of the DTR contigs were ≤15kb, which is a plausible size for complete RNA virus genomes. A more detailed gene annotation of the 101 representatives contigs for these DTR sequences affiliated to *Riboviria* revealed 3 main groups. First, 68 contigs encoded an RdRP where the closest relative in NCBI Viral RefSeq was found within the Narna-like clade. Genomes from this RNA virus group, which includes mitoviruses, were previously observed to assemble as circular contig, likely either because of the existence of a circular form of the genome or because of a replication mechanism involving a concatemer intermediary [6, 7]. These contigs, which represent the majority of the set, thus likely represent genuine complete *Riboviria* genomes. Another set of 15 sequences lacked an RdRP or other clear taxonomic marker gene but shared similarity to uncharacterized genes in known *Riboviria* genomes. The last set of 18 DTR contigs could be identified as members of the CRESS-DNA group (i.e. ssDNA viruses), based on the presence of a replication-associated gene typical from this group. These sequences represent complete genomes but were mis-affiliated as *Riboviria* instead of CRESS-DNA and were therefore excluded from Figure 2B and Figure 2C.

#### Additional analysis of the 528 kb viral contig from Ace Lake in Antarctica

The IMG/VR contig (IMG contig ID: Ga0222679_1000001) was identified from an Ace lake, Antarctica sample (IMG taxon ID: 3300022858) and predicted as complete based on the presence of a 127-bp DTR. The terminal repeat did not contain any low complexity regions and occurred three times on the contig (twice at termini and one other time). The contig was classified as viral based on a VirFinder p-value of 0.010 and score of 0.92 as well as the presence of 35 CheckV viral markers of 601 total protein-coding genes. Manual annotation also revealed the presence of a phage-like terminase large subunit (TerL) and a major capsid protein, two hallmark genes of phages in the *Caudovirales* order. 19 CheckV microbial markers were found, but these were interspersed between viral genes and did not result in CheckV predicting any host regions. A self-alignment of the contig with blastn did not reveal any large duplicated regions beyond the 127-bp DTR.

To validate circularity, we first ran CheckV and obtained an estimated completeness of 100%. The completeness estimate was based on a 100% ANI / 99.8% AF match to a CheckV sequence (DTR_285855) that was derived from a different sample from the same lake (IMG taxon ID: 3300025697, IMG contig ID: Ga0208769_1000001). As further validation, we performed read mapping from the sample (sequencing project ID: 1166905) to the 528,258 bp circular contig in order to test whether any reads spanned the circular breakpoint. After mapping with Bowtie 2 [8]using default options, we discarded paired end reads with more than 2 mismatches and discarded reads mapped to the same strand. After these filters 107,332 reads were mapped to the contig with a median insert length of 311 bp and read length of 150 bp. Supporting the circularity, we identified 10 reads with an insert length of 528,046 bp that spanned nearly the entire contig; assuming these reads instead spanned the circular breakpoint, then their insert lengths would instead be 212 bp, which is plausible for this dataset.

While *Caudovirales* genomes are typically ~50kb, larger genomes of ~500kb have been reported [9]. Recently, a set of new large (≥200kb) phages were reported from metagenome assemblies from which 10 major clades were proposed [10]. Based on a TerL phylogeny, contig Ga0208769_1000001 seems to be a new virus related to one of these clades ("Biggiephage", Figure S3B). Several members of the Biggiephage clade encode CRISPR arrays [10], and similarly contig Ga0208769_1000001 encodes a Type I-C-like CRISPR array (Figure S3A). No host could be predicted for Ga0208769_1000001 as no significant match was identified between this contig and the IMG CRISPR spacer database. Similarly, no significant match was identified between the spacers encoded on contig Ga0208769_1000001 and other Ace Lake contigs, hence it is unclear at this stage which elements are targeted by this CRISPR array. Finally, contig Ga0208769_1000001 included an unusually high number of transposases (14) distributed throughout the sequence, which suggests that mobile genetic elements may play a role in the large size of this genome.


1. Ren, J., et al., *VirFinder: a novel k-mer based tool for identifying viral sequences from assembled metagenomic data.* Microbiome, 2017. **5**(1): p. 69.
2. Chen, I.A., et al., *IMG/M v.5.0: an integrated data management and comparative analysis system for microbial genomes and microbiomes.* Nucleic Acids Res, 2019. **47**(D1): p. D666-D677.
3. Zimmermann, L., et al., *A Completely Reimplemented MPI Bioinformatics Toolkit with a New HHpred Server at its Core.* J Mol Biol, 2018. **430**(15): p. 2237-2243.
4. Paez-Espino, D., et al., *IMG/VR v.2.0: an integrated data management and analysis system for cultivated and environmental viral genomes.* Nucleic Acids Res, 2019. **47**(D1): p. D678-D686.
5. Gregory, A.C., et al., *Marine DNA Viral Macro- and Microdiversity from Pole to Pole.* Cell, 2019. **177**(5): p. 1109-1123 e14.
6. Bruenn, J.A., B.E. Warner, and P. Yerramsetty, *Widespread mitovirus sequences in plant genomes.* PeerJ, 2015. **3**: p. e876.
7. Hintz, W.E., et al., *Two novel mitoviruses from a Canadian isolate of the Dutch elm pathogen Ophiostoma novo-ulmi (93-1224).* Virol J, 2013. **10**: p. 252.
8. Langmead, B. and S.L. Salzberg, *Fast gapped-read alignment with Bowtie 2.* Nat Methods, 2012. **9**(4): p. 357-9.

9. Yuan, Y. and M. Gao, *Jumbo Bacteriophages: An Overview.* Front Microbiol, 2017. **8**: p. 403.
10. Al-Shayeb, B., et al., *Clades of huge phages from across Earth's ecosystems.* Nature, 2020. **578**(7795): p. 425-431.
